# Inferring reaction network structure from single-cell, multiplex data, using toric systems theory

**DOI:** 10.1101/731018

**Authors:** Shu Wang, Jia-Ren Lin, Eduardo D. Sontag, Peter K. Sorger

## Abstract

The goal of many single-cell studies on eukaryotic cells is to gain insight into the biochemical reactions that control cell fate and state. In this paper we introduce the concept of effective stoichiometric space (ESS) to guide the reconstruction of biochemical networks from multiplexed, fixed time-point, single-cell data. In contrast to methods based solely on statistical models of data, the ESS method leverages the power of the geometric theory of toric varieties to begin unraveling the structure of chemical reaction networks (CRN). This application of toric theory enables a data-driven mapping of covariance relationships in single cell measurements into stoichiometric information, one in which each cell subpopulation has its associated ESS interpreted in terms of CRN theory. In the development of ESS we reframe certain aspects of the theory of CRN to better match data analysis. As an application of our approach we process cytomery- and image-based single-cell datasets and identify differences in cells treated with kinase inhibitors. Our approach is directly applicable to data acquired using readily accessible experimental methods such as Fluorescence Activated Cell Sorting (FACS) and multiplex immunofluorescence.

**Author summary:** We introduce a new notion, which we call the effective stoichiometric space (ESS), that elucidates network structure from the covariances of single-cell multiplexed data. The ESS approach differs from methods that are based on purely statistical models of data: it allows a completely new and data-driven translation of the theory of toric varieties in geometry and specifically their role in chemical reaction networks (CRN). In the process, we reframe certain aspects of the theory of CRN. As illustrations of our approach, we find stoichiometry in different single-cell datasets, and pinpoint dose-dependence of network perturbations in drug-treated cells.

## Introduction

Single-cell, multiplexed datasets have become prevalent [1, 2], and include data on transcript levels measured by sc-RNAseq [3], protein levels measured by flow cytometry [4], or cell morphology and protein localization measured by multiplex imaging [5–8]. An obvious advantage of such data is that it makes possible the detection and quantification of differences among cells in a population, including those arising from cyclic processes such as cell division and differentiation programs that are asynchronous between cells [9,10]. A more subtle advantage of single-cell data is that they report on relationships among measured features, phosphorylation state of receptors and nuclear localization of transcription factors for example. Because such features are subject to natural stochastic fluctuation across a population of cells [11], measuring the extent of correlation between otherwise independently fluctuating features makes it possible to infer the topologies of biological networks [12,13].

A wide variety of tools have been developed for visualization of single cell data, including t-SNE [14] and MAPPER [15], and for generating networks from such data using Bayesian Networks [16] and machine learning [17]. In many cases, the goal of such tools is to produce statistical models. In this paper we describe an alternative analytical framework founded on reaction theory. We make the assumption that proteins in a compartment react with each other in a manner that is well approximated by the continuum assumptions of Mass-Action Kinetics (MAK) [18], the foundation of familiar biochemical treatment of reactions such as Michaelis-Menten kinetics and Hill functions [19–21]. Compartments in this formalism can be different macromolecule assemblies or different locations in a cell. Cellular biochemistry is complex, involving thousands of proteins and an unknown number of reaction compartments. Constructing dynamical systems of cellular processes based on MAK is computationally challenging, despite its theoretical appeal and analytical tractability. High-dimensional whole-cell dynamical models also suffer from a sparsity of data able to constrain such a model (although insights have been found by this approach [22, 23]).

Unexpectedly, we have been able to sidestep some of the challenges posed by MAK in a cellular context by leveraging geometric aspects of these dynamical systems and thereby obtain analytical conclusions from single cell data. Chemical Reaction Network Theory (CRNT) is a branch of dynamical systems analysis that relies primarily on topological features of the reaction network [24–26]. In this paper we frame results from CRNT in the context of multiplexed single cell data. We demonstrate that unexpected insights into the topologies of reaction networks can be derived from such data based on familiar and simple MAK principles. Specifically, from multiplexed flow cytometry (FACS) and multiplexed immunofluorescence (CyCIF) data, we observe integer stoichiometry of reactions, and show that four antimitogenic drugs perturb a cell’s reaction network in a largely dose-independent manner.

To illustrate this approach, we briefly review some basic definitions. For a reaction

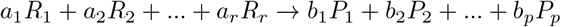

with reactants {*R_i_*}, products {*P_j_*}, and stoichiometric coefficients {*a_i_*} and {*b_j_*}, we associate a *reaction vector* 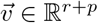, given by:

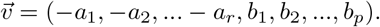

Provided the reverse reaction exists, the *steady state* concentrations of the reactions obey:

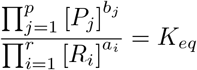

for some equilibrium constant *K_eq_*. This equality can be rewritten in terms of 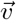, for the chemical concentrations 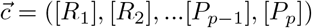:

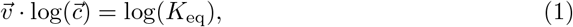

where the logarithm of the vector is defined as the element-wise logarithm. (Logarithms are taken in any fixed basis, for example decimal.) Observe that this is a linear equation on the reaction vectors, if one knows the (logarithms of) concentrations.

Given a network *G* composed of such reactions, the overall dynamics are described by a system of differential equations, in which the rate of change of any chemical species’ concentration is given by the sum of reaction rates in which it is a product, minus the sum of reaction rates in which it is a reactant [18]. As a simple example, consider a system of reactions:

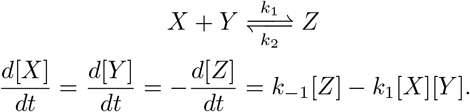

We will focus on two objects associated to such systems: the *steady state set E*, which is defined as the set of concentrations for which all the time derivatives vanish, and the *stoichiometric subspace S*, defined as the linear span of all the reaction vectors. In the example described above, the steady state set is a nonlinear surface, shown in Fig 1a, and its one-dimensional stoichiometric subspace is represented by a yellow line. The surface characterizes the network well, since any initial concentration ([*X*], [*Y*], [*Z*]) (represented by red dots), approaches the steady state set. In our studies, *E* will be determined from experimental data, and we will be interested in reconstructing *S*, or parts of *S*.

**Fig 1.**
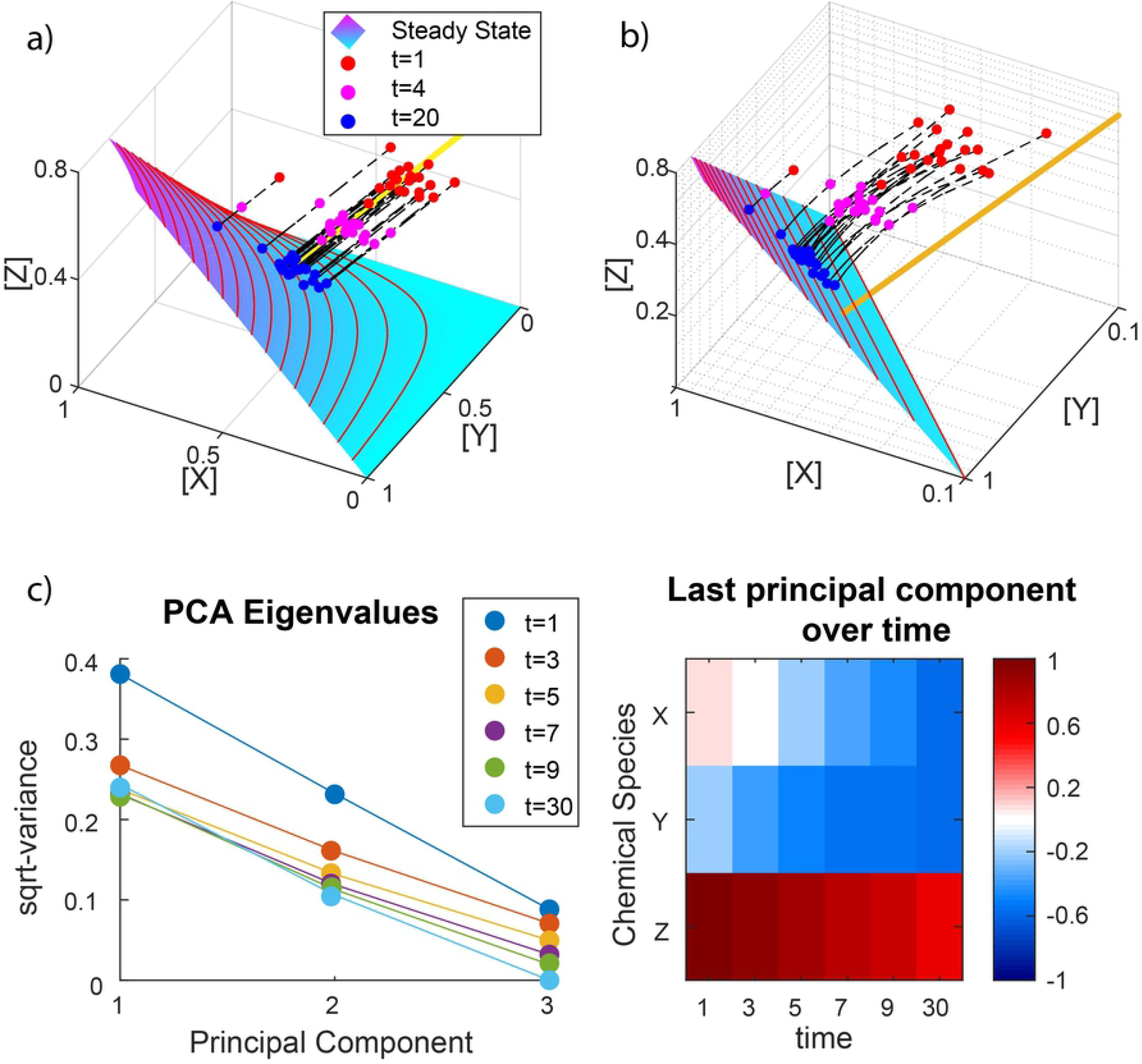
MAK dynamical systems and covariance in single-cell-data. (a) Several simulated trajectories of the reaction network *X* + *Y* ⇌ *Z* are shown. The steady state is shown in cyan/magenta, along with some of its level sets for fixed values of [*Z*]. A particular parallel translate of the stoichiometric subspace (coset) is shown as a yellow line. (b) Steady state in logarithmic coordinates. The orthogonal complement of this subspace (orange) is parallel to the stoichiometric subspace. (c) In log-concentration space, the covariance matrix of the chemical trajectories will have a decreasing eigenvalue for *t* → ∞, as evaluated by PCA, and the corresponding eigenvector will converge to the orthogonal complement, which is parallel to the stoichiometric subspace spanned by (−1,−1,1).

Among MAK dynamical systems, the subset known as “complex-balanced” reaction networks (which includes the familiar case of “detailed-balanced networks” [27]), has steady state sets that are easily expressed in terms of the stoichiometric subspaces [24]. Complex balancing means that each “complex” (a node of the reaction network, such as “*X* + *Y*” and “*Z*” in our example) is balanced with respect to inflow and outflow, analogous to a Kirschoff current law (in-flux = out-flux, at each node). It is a nontrivial fact that, for every 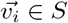, the steady state set *E* is precisely the set of all those vectors 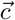 that satisfy all the following equalities:

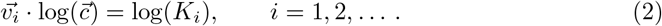

This is analogous to the case of a single reaction in Eq 1, except that *K_i_* is not the equilibrium constant of the isolated reaction, but is instead a constant that accounts for kinetic constants from the entire network. These equalities imply that, in log-concentration space, the transformed steady state set, log(*E*) ≡ *V*, is an affine (linear with shift) subspace whose orthogonal complement coincides with *S*. Our earlier example was complex-balanced, so after taking the logarithm, its steady state surface becomes a plane in Fig 1b, whose orthogonal complement, in orange, is parallel to the yellow line, shown in Fig 1a. As another example of a complex-balanced reaction network, consider a network with reversible and irreversible reactions:

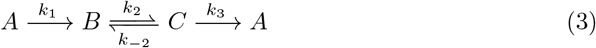

that has a steady state set satisfying (see S1 Appendix):

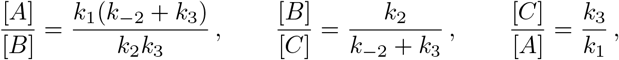

which are log-linear relations in the form of Eq 2. Although the above two examples are always complex-balanced, any network of any size can be complex-balanced if the kinetic constants *k_i_* are additionally constrained [25].

More generally, the subset of reaction networks that obey log-linearity are called *toric* in the algebraic-geometric CRNT literature [25,28]. This log-linearity greatly simplifies the analysis of a nonlinear problem, which is the key appeal in our making a MAK assumption. The current work is concerned with such *toric* systems, of which complex-balanced reaction networks are the best-known example.

## Results

### Overview of the Approach

We represent a single cell by a vector that includes as components the concentrations 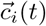 of relevant chemical species. We assume that all cells in the population being studied are governed by a common, complex-balanced, MAK reaction network *G* with reaction constants {*k_G_*}. The localization of a reactant into different cellular compartments (e.g. nucleus and cytoplasm) or different macro-molecular complexes is managed using the conventional compartmentalized formalism and simply adds elements to 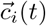. As we will see, the fact that *G* is ineffably complex does not limit our theoretical analysis.

We reframe the equations described in the introduction in terms of the distribution of chemical trajectories from a population of cells, 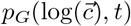, making it possible to approximate the stoichiometric subspace *S* of *G* from a fixed-time sample distribution 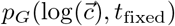, where *t*_fixed_ is large in an appropriate sense. Typically, it is only possible to observe a subset of the species in a cellular reaction network. We find that when only a subset of the chemical species are observed, 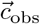, the covariance of 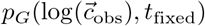 still makes it possible to determine a subset of *S*.

Exploring non-complex-balanced networks by simulation and examples, we find that our analysis method still recovers subspaces tied to network topology, analogous to how S is tied to reaction vectors. The key extension is that certain reaction networks that are non-complex-balanced can still have steady state set contained in a toric manifold (either exactly or approximately), whose orthogonal complement in log-concentration space has a straightforward relation to network topology.

With this theoretical background, we show that single-cell, multiplexed data (sc-data) that can feasibly be obtained from mammalian cells using multiplexed flow cytometry (FACS) or multiplexed immunofluorescence (by CyCIF and other similar methods) can be effectively analyzed within our framework, just using MAK assumptions. In particular, we find that (i) Principal Components Analysis (PCA) of single cell data produces principal components (PCs) that lie on near-integer subspaces, which our framework interprets as the stoichiometric constants in the underlying reaction, and (ii) for cells exposed to different small molecule inhibitors of regulatory proteins (primarily protein kinase inhibitors), the covariance structure is conserved over a range of concentrations for any inhibitor, which our framework explains as the conservation of reaction network topology.

### Single-cell covariance from complex-balanced reaction networks

Suppose that a population of *N* chemical trajectories 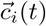 is governed by a complex-balanced, MAK reaction network *G*, with stoichiometric subspace *S* and steady state set *E*. As *t* → ∞, 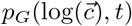 approaches a distribution supported on *V* ≡ log(*E*), whose sample covariance matrix Σ is a singular matrix with singular eigenspace equal to *S* (see Methods).

Applied to the example in Fig 1, the trajectories of *X, Y*, and *Z* concentrations at any time t constitute a dataset whose sample covariance matrix has one eigenvalue approaching zero as *t* → ∞ (Fig 1c). This eigenvalue’s corresponding eigenvector approaches (−1,−1,1), whose span is the stoichiometric subspace represented earlier by the orange line in Fig 1b.

In general, if we identify each cell in a population with a vector for the concentrations of all its relevant biochemical species 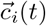, the hypotheses above allow us to extract the stoichiometric subspace *S* of the underlying reaction network by eigendecomposition of the sc-data covariance. This computation is commonly performed by PCA [29]. Whereas the literature typically considers the PCs that explain the greatest variance (e.g. PC1-3), we are interested in the singular eigenspace *S*, which is spanned approximately by the principal components that explain the least variance. To identify *S* with real data we look for a gap in the eigenvalue spectrum: the eigenvalues converging to 0 will be small and similar in magnitude, forming a cluster, while the remaining, larger eigenvalues will appear separate from that cluster. When such eigenvalues are arranged in ascending order, a gap appears right after the last eigenvalue of the small cluster. Such a gap is unexpected under the null hypothesis that the data is drawn from a random, multivariate normal distribution with equal variance in all directions [30]. Finding a gap in real data is nontrivial, and we discuss this subtlety in later sections where we analyzed FACS data.

### Timescale separation

For finite but sufficiently long times *t*, information about timescales can be found in sc-data. The eigenvalue spectrum of Σ, under the hypotheses described above, has at least one “gap” - a region of nonuniform spacing between neighboring eigenvalues - which separates the eigenvalues into “small” and “large” values. The small eigenspace approaches *S* as *t* increases. Additional gaps may indicate embedded subspaces *S_i_* ⊂ *S*, spanned by the reactions that occur on faster timescales, so that we have *S*_1_ ⊂ *S*_2_ ⊂ … ⊂ *S* corresponding to different cutoffs for “fast” and “slow” (see Methods).

Following our earlier example of 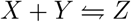 in Fig 1, we add a reaction 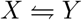 whose kinetic constants are substantially slower. The trajectories now converge first to the earlier surface, since it is the steady state of the fast reaction. With enough time, those trajectories eventually converge to the steady state of both reactions (see Fig 2a), which is now a curve embedded in the surface. This separation of timescales is studied formally using *singular perturbation theory* [31] for dynamical systems, in which the first surface is the *slow manifold*, because trajectories converge quickly to its neighborhood, before undergoing slow dynamics constrained to that neighborhood.

**Fig 2.**
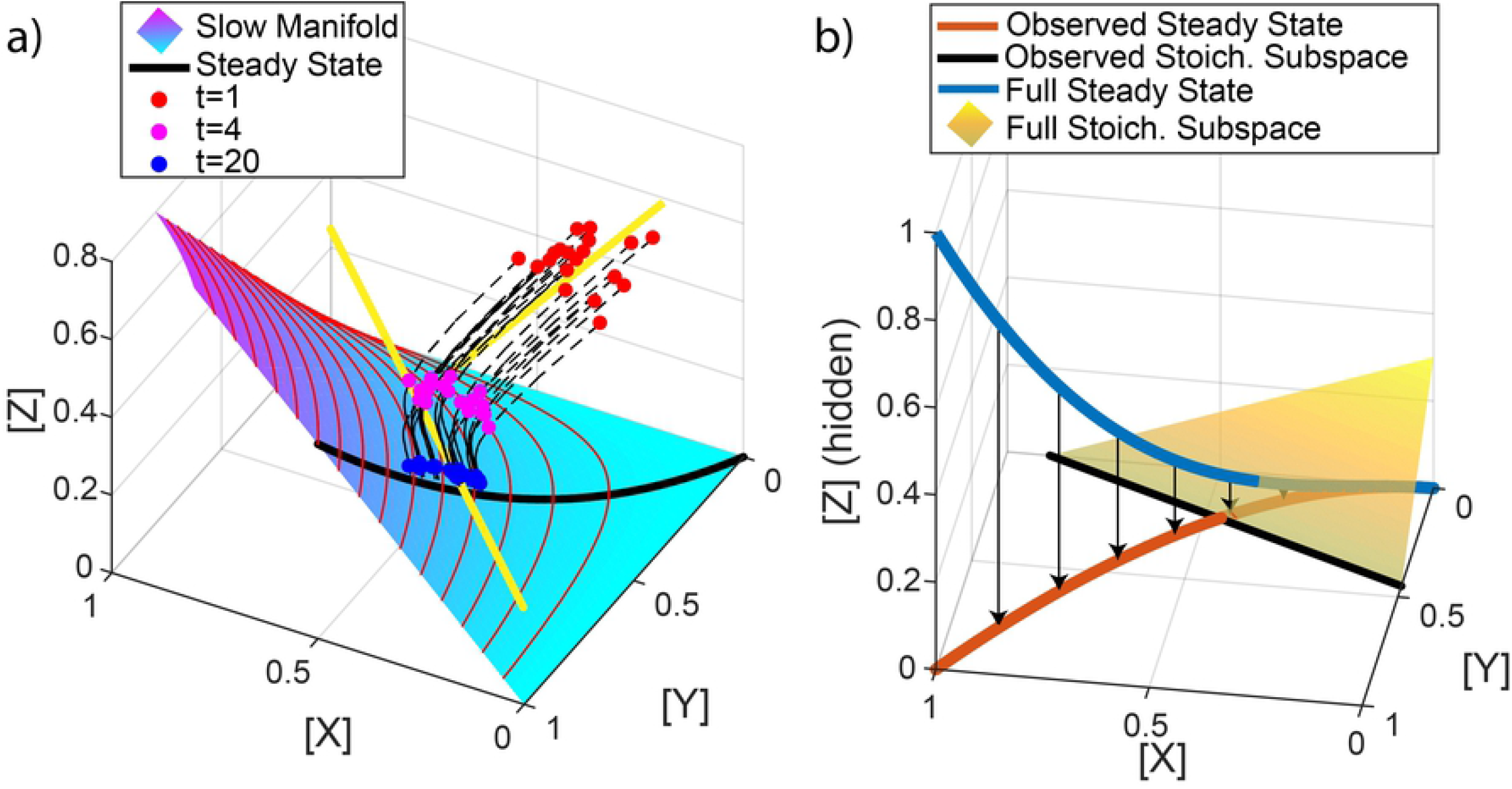
Timescale separation and hidden variables. (a) Simulated trajectories are shown for the reaction network with the additional, slow reaction 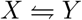. Trajectories first converge toward the steady state set of the fast reactions alone, the slow manifold, before slowly converging to the complete steady state (black). (b) An example of a log-linear steady state set (blue) and its stoichiometric subspace (yellow) are depicted. Supposing we observe *X* and *Y*, but *Z* remains hidden, we see the projected steady state set (orange), which is still log-linear. The orthogonal complement we would observe in log-concentration space is the intersection (black) of the original stoichiometric subspace and the observable plane.

For detailed-balanced reaction networks, slow manifolds are approximately the steady state sets of fast networks defined by ignoring slow reactions [32], and one might expect this to be true more generally. If so, then just as a single gap appears when trajectories converge to the full network’s steady state manifold, another gap appears as trajectories converge to the fast network’s steady state manifold. The larger the timescale separation, the larger the gap. Since there can be many separated timescales in a network, we expect correspondingly many gaps. Of note, these gaps separate all the PCs into timescales, with the largest PCs’ span representing the infinitely slow timescale.

### Accounting for unobservables: net reactions

MAK assumes well-mixed, elementary reactions involving the collision of molecules, but single-cell experiments never provide data on all, or even most, of the chemical species participating in elementary reactions for any given biological process. However, assuming that MAK adequately describes the elementary reactions, our conclusions change minimally after accounting for these unobserved species, thanks to log-linearity. More specifically, given a complex-balanced MAK network *G* that describes the dynamics of *N* chemical species in ℝ^*N*^, with stoichiometric subspace *S* and steady state set *E*, suppose that only a subset *n* of the species in *N* is observable. The observed set *E*_obs_ is then the orthogonal projection of *E* onto ℝ^*n*^ ⊕ 0 ⊂ ℝ^*N*^, and is still a log-linear set. The orthogonal complement *S*_obs_ of *V*_obs_ ≡ log(*E*_obs_) is precisely:

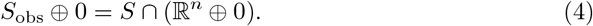

That is, *S*_obs_ is the intersection of the stoichiometric subspace and the observable space (see Methods).

As an example in Fig 2b, suppose *N* = 3 chemicals *X, Y, Z* obey MAK, but we only observe *n* = 2 of them, *X* and *Y*. If the steady state set *E* is a one-dimensional, log-linear curve, in orange, then *S* is a two-dimensional plane. Thus, in the observed ℝ^2^, we see the projection of *E, E*_obs_, in blue, whose orthogonal complement in log-concentration space, *S*_obs_ shown in black, is the intersection between the plane *S* and the observed plane ℝ^2^ ⊕ 0.

The fact that the observed orthogonal complement *S*_obs_ is a subset of *S* is important. It implies that any 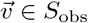 is a linear combination of the reaction vectors that span *S*. Intuitively, a linear combination of elementary reactions is a *net* reaction, just as glucose metabolism is often summarized by

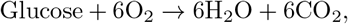

representing a sum of all the elementary reactions that occur during glycolysis and electron transport. As a further example, consider the earlier reaction in (3) and suppose that we only observe *A* and *B*. The steady state set *E* is a line through the origin in ℝ^3^, which is still a line after projecting into the observed ℝ^2^. The log-transform of any line through the origin in ℝ^2^ becomes a shifted line spanned by (1,1), whose orthogonal complement is spanned by (1, −1) (see S1 Appendix). Thus, by observing the projected line, and assuming complex-balancing, we can conclude that *A* ⇌ *B* is a net reaction in the full system, which indeed it is: the one direction is given by *A* → *B*, while the reverse direction is given by *B* → *C* → *A*.

In summary, not only is *S*_obs_ composed of net reaction vectors, the equality in Eq 4 of *S*_obs_ with the intersection implies that *S*_obs_ contains *every* net reaction that can be written in terms of the observed chemical species. In this sense, it is *maximal*.

### Networks other then those with complex balance may still have toric geometry

Whereas complex-balanced networks provide a sufficient condition for the previous results to hold, similar results hold for a larger class of MAK networks, relying on the log-linearity of steady states.

For example, take a simplified kinase(*E*)-phosphatase(*F*)-substrate(*S*) system:

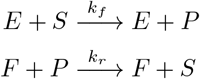

where the product *P* has steady-state:

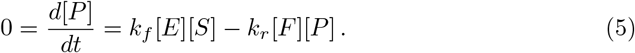

The complexes *E* + *S* and *F* + *P* are both reactant complexes, and they appear with opposite sign in the total rate of change of the species *P*. We therefore expect that the orthogonal complement should contain a vector denoting the difference between these complexes. By a quick rearrangement, we see that:

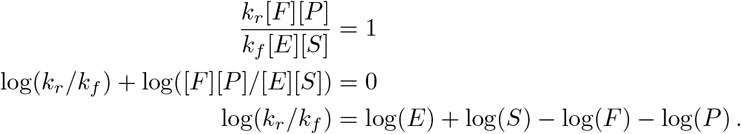

The orthogonal complement contains (1,1, −1, −1), which would be seen in data, informing us that *E* + *S* and *F* + *P* are reactant complexes that balance each other. The result is unchanged if we include the usual Michaelis-Menten enzyme-substrate complex, which is implicit in [28]. Thus, applying our method to data generated by a reaction network that has log-linear, or “toric”, steady states, the singular eigenspace still informs us about reaction topology.

Pérez-Millán et al. provide a sufficient condition for a reaction network to have “toric steady states” [28]. This broader class of networks even allows for multistability, which is strictly prohibited for complex-balanced networks. As in the example, the orthogonal complement *V*^⊥^ of steady states in log coordinates need not coincide with the stoichiometric subspace, although *V*^⊥^ still relates to network topology.

Furthermore, the steady state set need only be a *subset* of a log-linear set in order to extract the same information, although the set of reactions we recover is no longer maximal. Taking the previous example, add the reactions

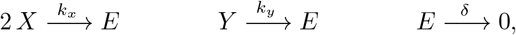

which imposes an additional, non-log-linear constraint on steady states, to the one in Eq 5:

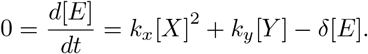

Despite this, the previous log-linear constraint in Eq 5 still ensures that at steady state, a sample of trajectories will have zero variance along (1,1, −1, −1) in the log-coordinates, as in the previous example.

In general, non-complex-balanced (and more generally non-toric) scenarios are not amenable to analytical treatment and we therefore explore them via simulation. As a reference, we first simulated complex-balanced reaction networks with 20 chemical species, including random single and binary reactions. Timescale differences were included by drawing the kinetic constants from two separate distributions. At different timepoints, the distribution of chemical trajectories was subjected to PCA (see Fig 3a). The eigenvalue spectra were found to exhibit gaps that grew larger with time. To confirm that the singular eigenspace spanned the defined stoichiometric subspace, we used Principal Angle Decomposition (PAD) to measure the difference in angles between the two subspaces [33]. We found that the angles converged to zero over time. The slower reactions led to distinctly larger eigenvalues, whose corresponding eigenvectors converged later. Such an example is shown in Fig 3a, where the network that was generated had a stoichiometric subspace of dimension 11, and the 11 PCs’ span converges to the subspace, as evaluated by principal angles. Some reactions were slower, leading to slower convergence along 2 dimensions, visible in the inset. This is accompanied by 2 of the 11 eigenvalues being distinctly larger than the rest, as expected from our previous discussion of how timescale separation manifests as differences in variance.

**Fig 3.**
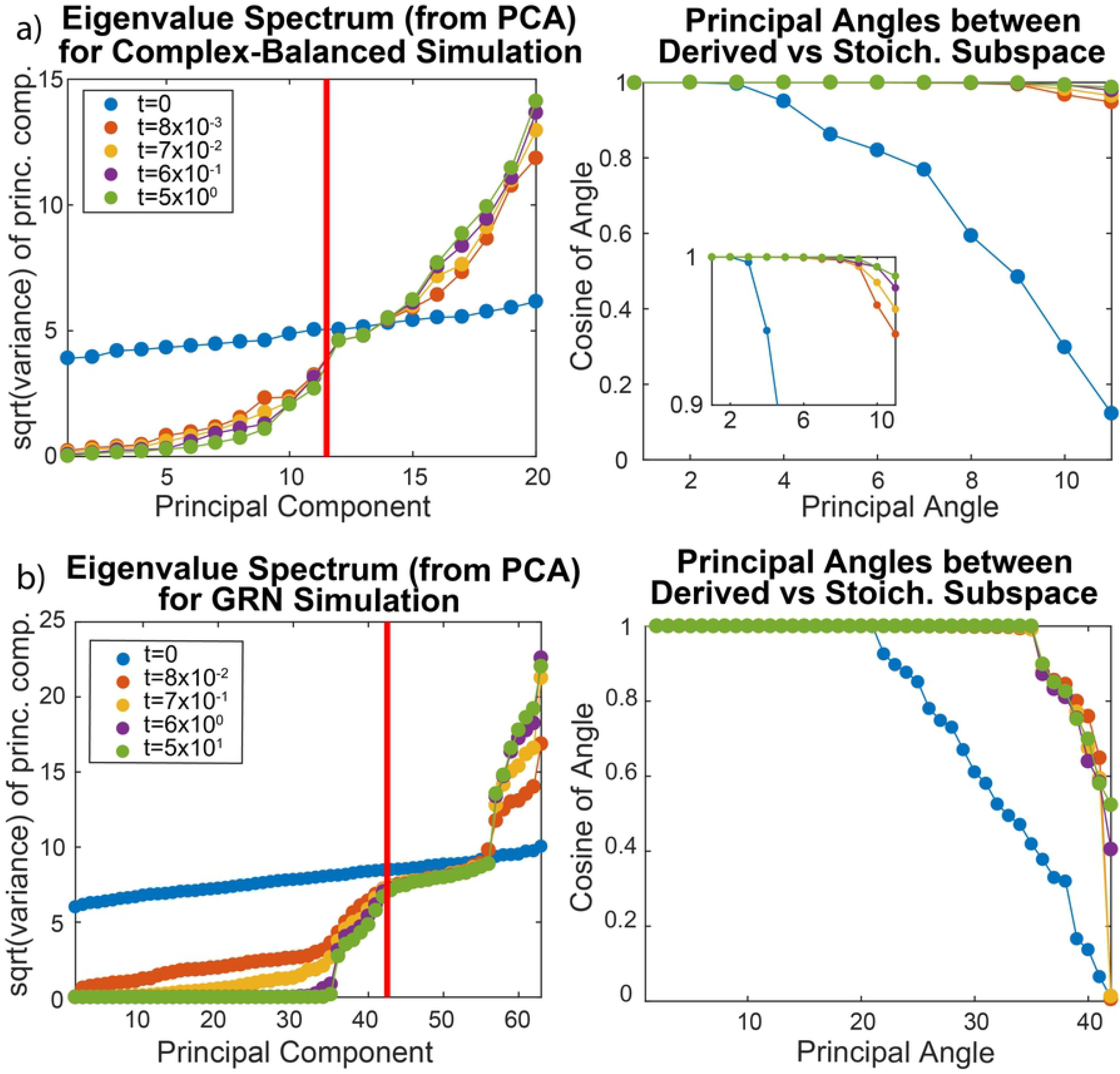
Reaction network simulations and deriving stoichiometric subspaces. (a) Example complex-balanced simulation, analyzed by PCA, shows 11 small eigenvalues, as expected from the simulated network’s structure, leading to a gap (red line) that grows larger with time. PAD shows that the span of these 11 eigenvectors converges to the true stoichiometric subspace. The 10th and 11th eigenvalues decrease slower than the others, due to slow reactions in the simulation. (b) An example GRN simulation for *n* = 7 is shown. From PCA, a gap in eigenvalues occurs at the expected dimension of the stoichiometric subspace (red line), as well as after the 35th eigenvalue. From PAD, the first 35 eigenvectors span the same subspace as the reversible binding reactions. The remaining 7 eigenvectors before the gap, whose eigenvalues are not as small, span a log-linear space tilted away from the stoichiometric subspace by angles between *π*/6 ~ *π*/3.

Having validated our conclusions about single-cell data covariance on a complex-balanced simulation, we turned to a non-complex-balanced model. We simulated a Gene Regulatory Network (GRN) with *n* genes *G_i_, n* corresponding to protein products *P_i_*, and ~ 70% of the possible *n*^2^ protein-bound genes 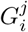 (*i*’th gene bound by the *j*’th protein) corresponding to proteins that function as transcription activators and repressors. The reactions in the network consisted of irreversible processes that resulted in protein production/degradation, and reversible binding of regulatory proteins to genes:

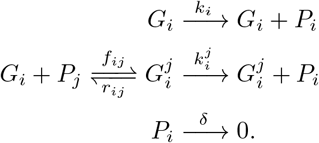

Transcription/translation was lumped into a single, protein production step for sake of simplicity. Analysis demonstrated that networks of this type are indeed non-complex-balanced (see Methods).

When the distribution of trajectories at the end of the simulations was analyzed by PCA, the eigenvalues of the covariance matrix for all simulations exhibited gaps visible in Fig 3b, indicating log-linear constraints. One gap always occurred after *d* eigenvalues, corresponding to the stoichiometric subspace’s dimension as computed symbolically. To evaluate the gap after *d* – *n* eigenvalues, we performed PAD on the first *d* – *n* eigenvectors and the stoichiometric subspace for the subnetwork of reversible binding reactions, finding that all principal angles were near 0. However, the remaining *n* eigenvectors converged to a subspace tilted *π*/6 ~ *π*/3 away from the stoichiometric subspace.

To understand the convergence of the reversible reactions and the n-dimensional tilt, consider a simple example of 2 genes’ concentrations, *g_A_,g_B_*, that of their protein products *p_A_,p_B_*, and one protein-bound gene, 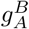. In this case the steady state equalities include:

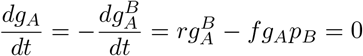

from which we retrieve the reaction vector for the reversible binding of protein *B* to gene *A*. Now, setting the rate of change of protein *B* to zero, by substitution we have:

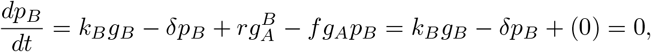

giving an orthogonal vector that connects *g_B_* and *p_B_* in a 1 and −1 ratio, even though this is not a reaction vector. Finally, we have

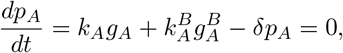

which does not give a log-linear relation. However, in various limiting cases, it is still possible to recover an asymptotically log-linear relation. For example, consider the common scenario in which protein *B* is an activator for gene *A*, so that 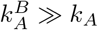:

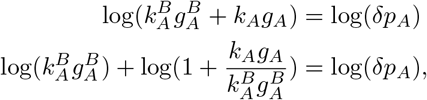

and for small 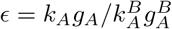, we recover a log-linear relation by a Taylor expansion of the middle term to zero’th order:

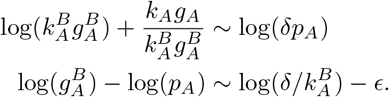

This possibly explains the origin of the *n* eigenvectors that were found to be tilted relative to the stoichiometric subspace: there are *n* such 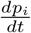 terms in the simulation, each giving an orthogonal vector ~ (1, −1) (the first coordinate being the *i^th^* protein species and the second being the most active bound-state of the *i^th^* gene), which is tilted *π*/4 from the *i^th^* protein’s reaction vector (1,0). In our simulation, multiple protein-bound variants existed for any gene, which adds *ϵ* error terms that may skew the angles further.

From this one small example, we see that log-linear constraints arise from complex-balanced reactions, from a balance between production and degradation, and from a biological, asymptotic case. We expect log-linear constraints to be mechanistically informative, even without complex-balancing, and thus our framework may be useful with further development in the analysis of general biological systems.

In the remainder of this paper we refer to the orthogonal complement of the minimal, linear set containing the log steady state as the *effective stoichiometric space* (ESS).

### Single cell data obtained by FACS has sparse covariance with integer structure

We analyzed a previously published multi-parameter Fluorescence-Activated Cell Sorting (FACS) dataset in which the levels of 11 phospho-proteins in the ERK/Akt signaling pathway were measured in primary human naive CD4+ T-cells [16]. Measurements were made in the presence of 14 different inhibiting or activating perturbations of the pathway. One of the conditions contained no signal for some phosphorylated species (probably for technical reasons), so we did not include the condition in our analysis.

FACS data from each condition were fit with a two-component Gaussian Mixture Model (GMM) to distinguish two empirical subpopulations, and the larger component was analyzed further. For each condition, the covariance matrix was eigendecomposed. Each eigenvalue spectrum showed at least one gap, denoted by an orange arrow in Fig 4a; in some cases an additional gap was visible, attributable to timescale separation.

**Fig 4.**
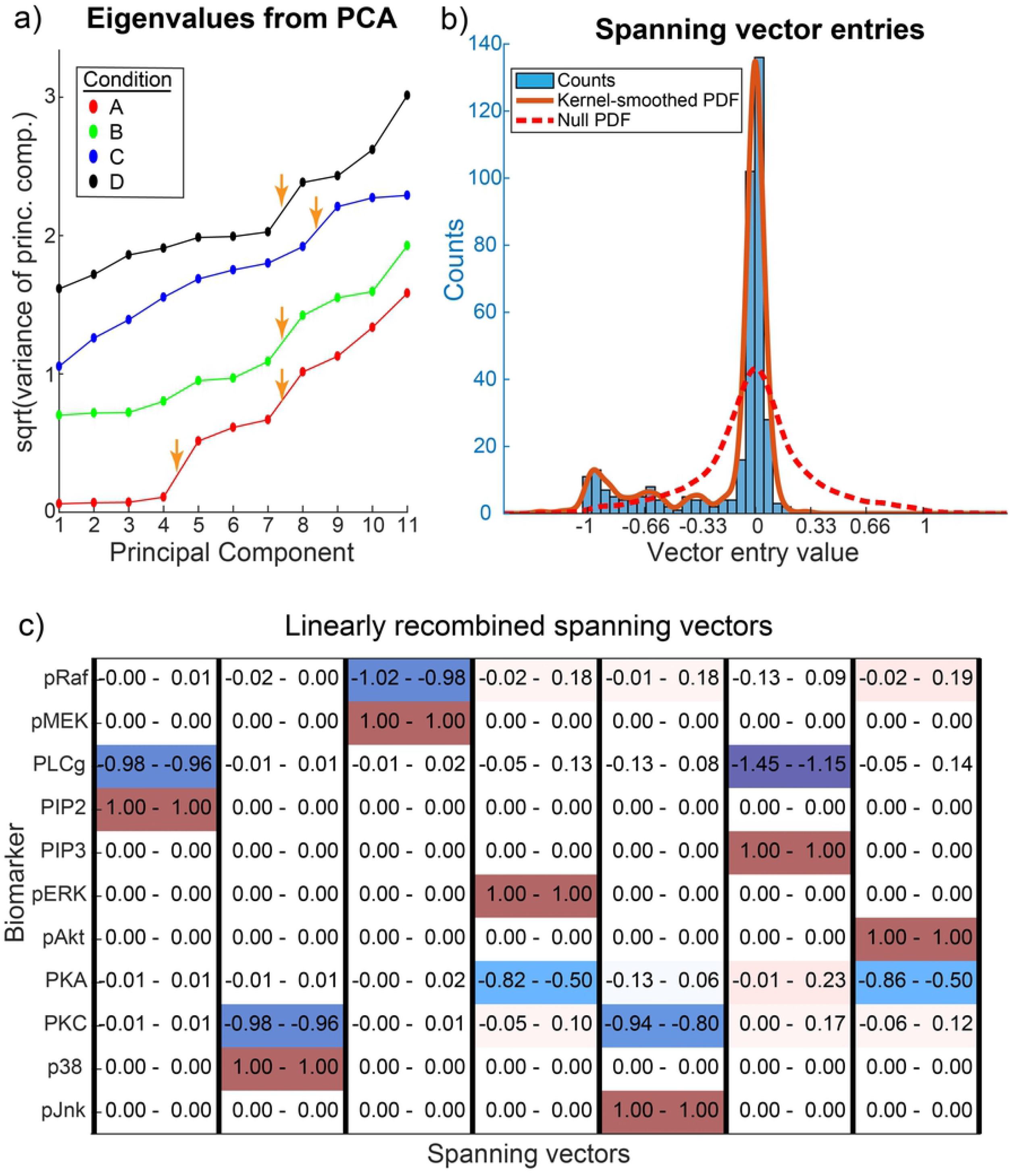
Peculiar properties of single-cell high-dimensional datasets (FACS) (a) Eigenvalue spectra from PCA of the dominant CD4+ subpopulations are shown for 4 of the 13 conditions (shifted to avoid overlap). Apparent gaps denoted by orange arrows. (b) The small eigenvectors were linearly recombined by row reduction on their transpose, with complete pivoting for ease of interpretation. The distribution of the linearly recombined entries from all 13 conditions are shown in a histogram (not including the 0s and 1s that are necessarily produced by row reduction), as well as with a Gaussian smoothing kernel of bandwidth 0.04. Peaks seem to appear at −1/3, −2/3, and −1. The null distribution for random, sparse, constraints is also shown for comparison. (c) As an example, the recombined vectors for Condition A are shown, with bootstrapped 95% confidence intervals. Other conditions are similar in appearance.

Gaps in eigenvalue spectra were identified by visual inspection, based on the presence of abrupt discontinuities, but the approach is not rigorous. Principled methods exist to identify which gaps are significant [30], but these methods apply only in the asymptotic limit when the number of dimensions *d* → ∞, with assumptions on the noise distribution. Thus multiple heuristic methods have been developed, such as looking for spikes in the slope of the spectrum, to choose component numbers in PCA; an overview and comparison of some methods are and given in [34,35]. In the current work we used such a heuristic approach to gap identification; the approach could potentially be improved with future research.

Each ESS was defined by choosing the gap farthest right. The corresponding eigenvectors were then interpreted by linearly recombining them by row reducing their transpose with complete pivoting [36]. This made it possible to represent the same linear subspace with sparser vectors whose entries are normalized to an arbitrary chemical species. The resulting vectors for a particular condition, are shown in Fig 4c, with a red entry with value 1 denoting the algorithm’s chosen normalizing species in each column. Each column can be interpreted as an effective, net reaction, in the broader sense. These data-derived ESS, for each condition, partitioned the 11 markers into groups, implying that steady-state values of chemical species stoichiometrically constrain each other only within these groups (see S1 Fig). The grouping from one condition was identical to that originally generated by Bayesian Causal Network Inference in Sachs et al. when applied to one condition (the specific condition is not identified in the manuscript, see [16] Supplement). While the method in [16] pooled the 14 perturbations to infer causal directions, our framework regards each perturbation as a change in the equilibrium constants and topology of the network, without imposing causal structure.

Additionally, the recombined eigenvectors’ entries (excluding the 1s and 0s necessarily produced by row reduction) had a distinct distribution (see Fig 4b). First, most entries were near zero, distributed between −0.2 and 0.2, which suggests a nontrivial sparseness in the span of the selected principal components. The asymmetry of the distribution is also unexpected (see Methods), but is a consequence of our framework: most reaction vectors involve both production and consumption, whose entries necessarily have opposite signs, so after eliminating the positive values of 1 generated by the row reduction algorithm, the remaining nonzero entries of those vectors should always include negative entries. This left 83 entries smaller than −0.2 that our framework expects to be small-integer-ratios, since reactions typically have small-integer stoichiometry. We notice that the entries bias towards 1/3, 2/3, and 1. These features are significantly different from the null expectation of a random and sparse structure underlying the data, even when accounting for how we heuristically chose the gaps (See S2 Appendix).

### CyCIF data covariance is also sparse and integer-like

We also analyzed a Cyclic Immunofluorescence (CyCIF) dataset that comprises measurement of the levels and modification states of 26 antigens with a focus on phospho-states of proteins involved in apoptosis, Akt/Erk signaling, cell cycle progression, and cytoskeletal structure. The dataset is found in the Library of Integrated Network-based Cellular Signatures (http://lincs.hms.harvard.edu/db/datasets/20267/). Nontransformed MCF10A mammary epithelial cells were exposed to four different kinase inhibitors at six doses each, totaling 24 distinct conditions. Data from each treatment were fit by a GMM with 1 ~ 4 components, where one component was always substantially larger than the rest; we refer to this component as *dominant*, and focused our analysis on it.

The eigenvalue spectra for each of the treatments also exhibited gaps, as denoted by orange arrows in Fig 5a. Defining each ESS using the gaps around the 10th-14th eigenvalues, row reduction of the selected eigenvectors once again generated the sparse, asymmetric distribution of vector entries observed for FACS data, with a bias to integer-ratios of −1, and possibly −1/2 (see Fig 5b), although less clear than in the FACS case. For each condition, the row reduced vectors suggest net reactions that sensibly relate the various proteins. For example, in one condition shown in 5c, total S6 was linked with mTor, and phosphorylated S6 at site S235 was linked to phospho-S6 at site S240, which matches the canonical picture that these proteins influence one another in the mTor-S6 signaling cascade. However, E-Cadherin’s contribution to the vector linking S6 with mTor, and the vector linking gamma-H2AX with PCNA, are less expected. The former may reflect the effect of mTor on the Epithelial-mesenchymal transition (EMT) [37], and the latter may reflect the involvement of S phase (as scored by PCNA) and gamma-H2AX in DNA repair. These biological details will require further analysis but the key point is that single cell microscopy (CyCIF) data resembles FACs data with respect to sparsity, integer-ratio entries, and the appearance of sensible connections between sets of proteins.

**Fig 5.**
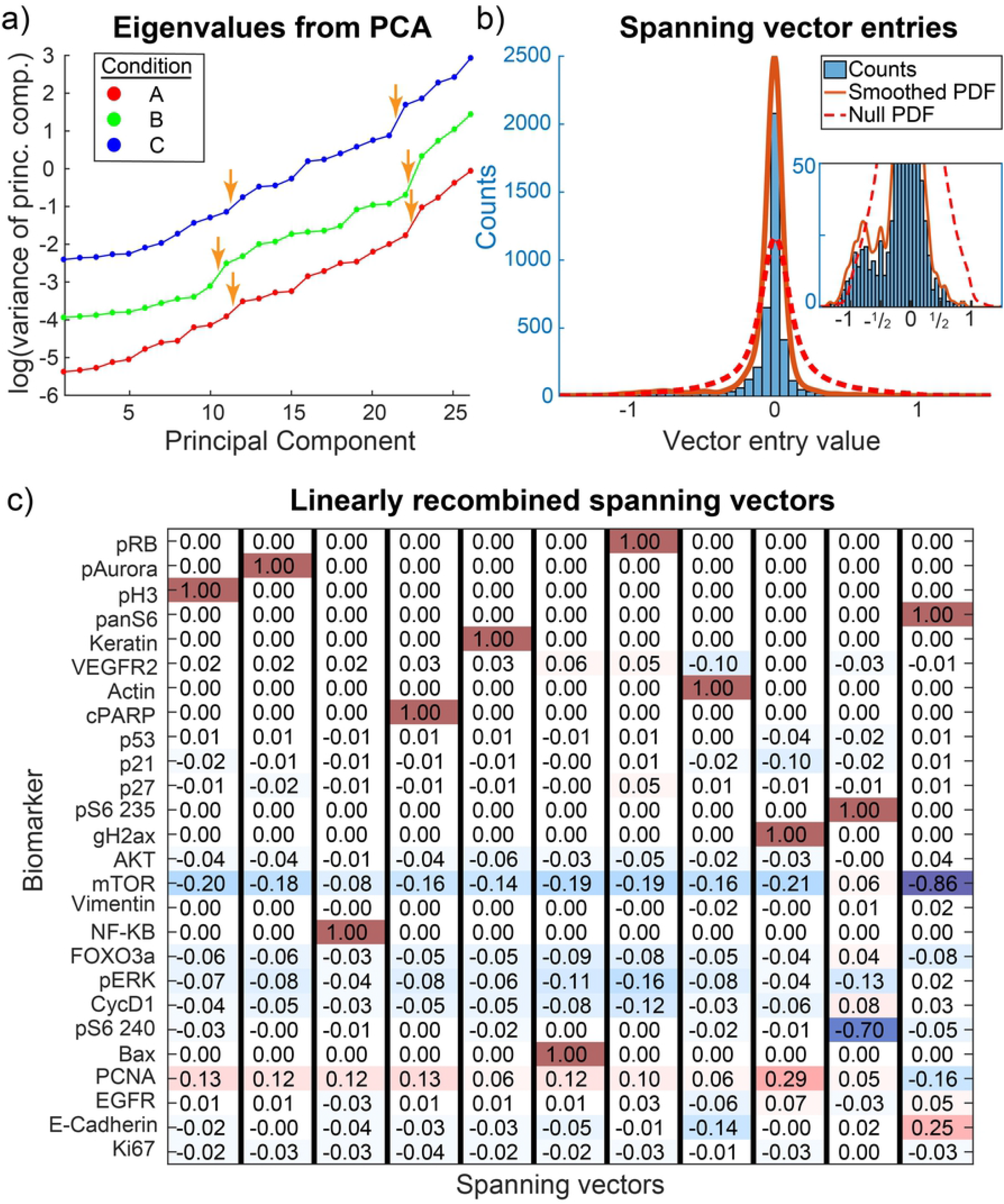
Peculiar properties of single-cell high-dimensional datasets (CyCIF) (a) The dominant subpopulations of the MCF10A cells were analyzed by PCA and the eigenvalue spectra are shown for 3 of the 24 conditions (shifted to avoid overlap). Some, but not all, apparent gaps denoted by orange arrows. (b) Singular eigenvectors were linearly recombined by row reduction on their transpose, with complete pivoting. The distribution of entries is displayed, along with a null. (c) Condition A’s dominant subpopulation’s recombined, singular eigenvectors are shown. Net reactions link the various proteins, such as S6 with mTor, or the two phosphoforms of S6 (235 and 240). Other conditions show similar sparseness.

### Data from drug-treated cells conserve covariance structure over large dose ranges

For the CyCIF data, we analyzed the dose and drug dependence of the ESS associated with the dominant fluorescent signal in each channel. To compare the subspaces from any two treatments, we first performed PAD between all pairs of treatments, and then summarized the principal angles *θ_i_* with the metric

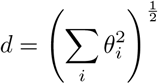

that appeared in [38]. To interpret this metric, the subspaces being compared must have the same dimension. Thus, we could not use our previously chosen gaps in the eigenvalue spectrum to define ESS for comparison between two treatments; instead, we chose the first 10 eigenvectors in each treatment condition to span a rough ESS for inter-treatment comparisons.

Between with and without drug (DMSO-only control samples), the ESS changed substantially, as shown in Fig 6a. This is expected, since the addition of a kinase inhibitor alters the set of reactions in the network, directly changing the ESS. As a simple example, consider the kinase-phosphatase-reaction system from earlier as part of a signaling cascade, but add an inhibitor for S’s downstream enzymatic activity, *D*:

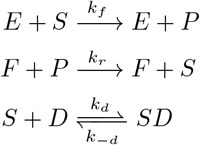

which leaves *d*[*P*]/*dt* unchanged, so (1,1,−1,−1) is still in the ESS. However, *d*[*S*]/*dt* ≠ *d*[*P*]/*dt* after the addition of *D* at concentration [*D*_0_]:

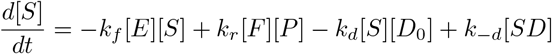

**Fig 6.**
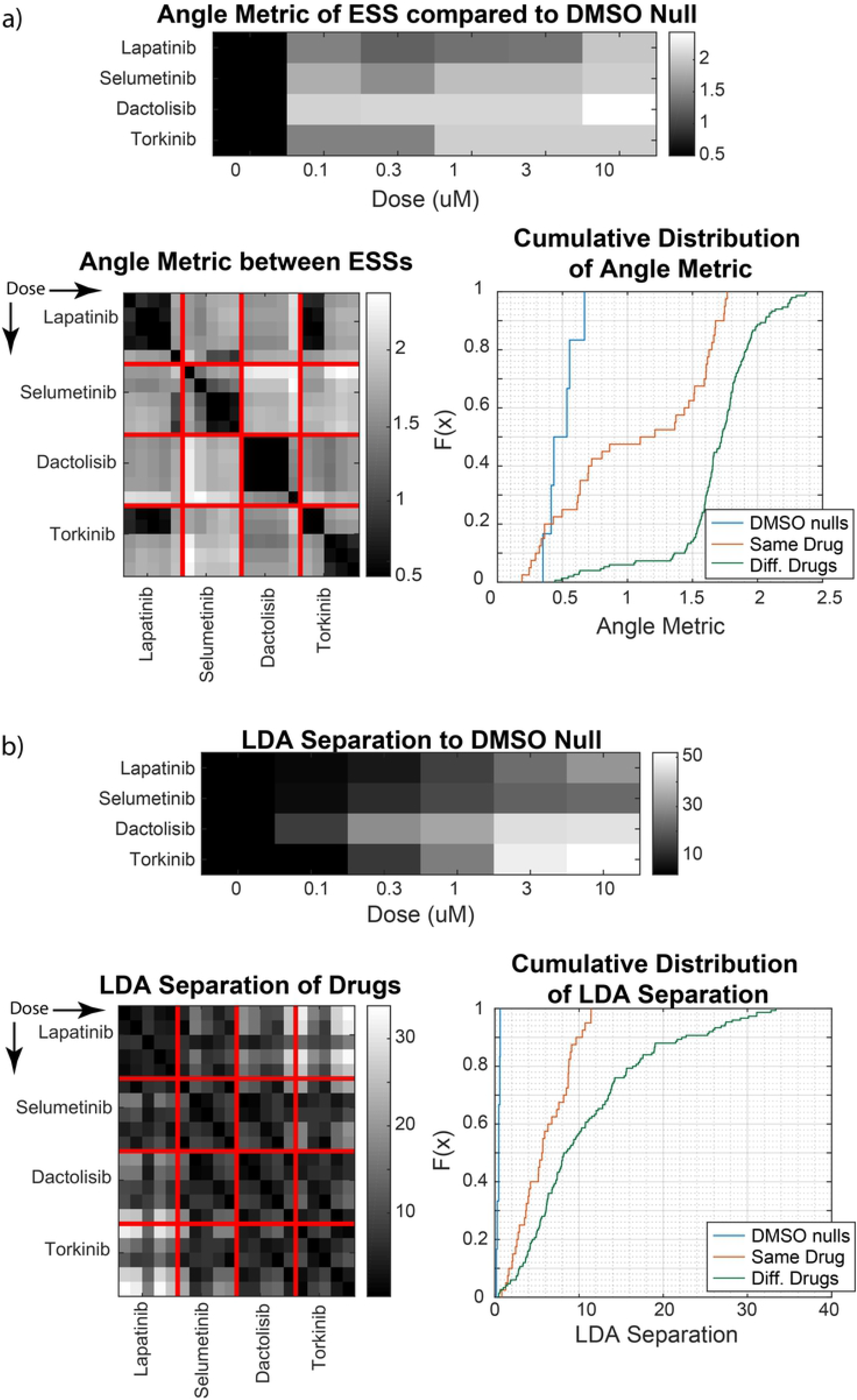
Comparison of reaction networks between drug treatments. (a) Analyses of the ESS between conditions, as quantified by an angle-based metric for a common cutoff of a 10-dimensional ESS. Average metric between a drug treatment and the four DMSO replicates (top) are plotted, as well as between any pair of non-zero doses of drug (bottom), arranged by increasing dose. The cumulative distribution of the metric is shown for the pairs between DMSO null replicates, the pairs that used the same drug, and the pairs that used different drugs. (b) Analyses of the LDA separation between the high-dimensional marker distributions, with analogous comparisons as above.

In various limits, we would expect new log-linear constraints. For example, if *k_d_*[*D*_0_] ≫ *k_f_*[*E*] and *k_r_* [*F*][*P*] ≫ *k*_−*d*_[*SD*], we observe the log-linear constraint (0,1,−1,−1).

Between treatments, the data shows the ESS remaining independent of dose [Do] almost 50% of the time with a precision comparable to experimental error (based on comparisons between the DMSO-only control samples), as shown in the cumulative distribution of Fig 6a. Between drugs, the ESS differed substantially: ~ 95% of comparisons showed a larger difference than the null. This is consistent with the two ideas: 1) changes in the dose of an inhibitory drug modify kinetic constants (phosphorylation rates), which is only expected to change the ESS in asymptotic limits, and 2) different drugs interact with different components of the network, resulting in different ESS. The dissimilarities of ESS that do occur between the same drug, do so between dose regimes as opposed to randomly, which is consistent with dose-dependent asymptotics. In the case of Torkinib, this may correspond to its reported polypharmacology [39]. Meanwhile, low-dose Lapatinib and low-dose Torkinib show similar ESS, as do high-dose Lapatinib and high-dose Selumetinib. This implies that in these corresponding dose-regimes, the drugs have similar topological effects on the 26 observed biomarkers, which is plausible given that Torkinib’s target (mTOR) and Selumetinib’s target (MEK) are in two pathways downstream of EGFR, which is Lapatinib’s target. These biological interpretations of the data remain preliminary, but the ESS has allowed us to pinpoint topological changes in reaction network stoichiometry, independent of parameter changes.

As a more conventional analysis of population differences, we also compared treatments using the Linear Discriminant Analysis (LDA) [40] separation between the dominant Gaussian components of any pair of conditions. The results are shown in Fig 6b, using the same format as the ESS angle-metric comparisons. Almost no pairs have a separation of comparable magnitude to experimental noise, and the magnitude of difference within the same drug is comparable to that between different drugs. Therefore, using LDA, we cannot discern the topological similarity between treatments that was obvious using ESS.

However, the LDA results do tell us that between drug treatments, the mean marker expression of the dominant populations change substantially. Changes in the mean correspond to changes in the equilibrium constants *K_i_* from Eq 2. Thus, the LDA result showing dose-dependent shifts, combined with the result of dose-independent ESS, allows for a rigorous interpretation: changing drug dose induces parallel shifts of the high-dimensional distribution of cells in marker-space, and changing the drug induces tilts of the distribution.

## Discussion

Single-cell, multiplex imaging and flow cytometry are increasingly used to identify cell states and study regulatory mechanisms. A range of computational methods have been developed to analyze the resulting high-dimensional data but most approaches are statistical. In this paper we explore the possibility of using insights from algebraic-geometry developed for Chemical Reaction Network Theory (CRNT) in the analysis of single-cell data. We find that an effective stoichiometric space (ESS) can be generated from such data to guide reconstruction of biochemical networks. In an initial test of our approach, interpretable network features were obtained from both synthetic and real experimental data. The advantage of using CRNT in this setting is that it provides a principled way to incorporate fundamental knowledge about how biomolecules interact through time and space.

A characteristic of sc-data is that measured features (typically the levels, localization and modification states of genes and proteins) co-vary in individuals cells. In the face of random fluctuation, patterns of covariance potentially contain information on interactions between biochemical species. A key questions is how this covariance information should be analyzed to obtain insight into the underlying biochemical pathways. We find that eigendecomposition of covariance matrices from sc-data can be interpreted in terms of network stoichiometry and timescales, without model simulation, independent of kinetic parameters, and unhindered by unobserved species; the latter point is critical because most single-cell data is sparse with respect to the number of reactants than can be measured. These features of the ESS approach are a direct consequence of toric (log-linear) manifolds that arise from an assumption of mass-action kinetics, and hold even under the looser requirement that a steady state is a subset of an approximately toric manifold. In this case, the system does not need to reach a quasi-steady state (QSS) but the steady state set need only be asymptotically stable, and enough time should have passed for approach to this QSS. Thus, the results of toric analysis can be applied when a population of cells is approaching quasi-steady state on a relevant time scale, further expanding the situations in which the ESS framework is informative and applicable.

We tested our approach using synthetic data derived from various simplified reaction systems and also showed that it can be applied to both FACS and multiplex imaging datasets. We extract features from the data that are consistent with an interpretation in a reaction network framework: integer-like stoichiometries for interacting species, and independence of network topology on the dose of a single drug used to perturbed the network. Other kinds of sc-data, such as mass cytometry or sc-RNAseq, can potentially be analyzed using the same approach. Because this paper focuses on the theoretical aspects of toric geometries as applied to sc-data, we have not yet tested any of the biological conclusions derived from the analysis of real data. However, the interactions we infer are consistent with current understanding of well-studied human signal transduction networks and with previous publications [16]. More extensive single-cell experiments will be required to fully test the potential for ESS analysis to generate new biological insight.

Simulation of synthetic complex-balanced networks and GRNs suggests ways to tailor reaction network ODEs to better match sc-data. Assuming that the goal of fitting the network to data is to match the mean *μ* and covariance Σ of key analytes, our results show that it is possible to predict a partitioning of the eigenspace of Σ without actually simulating the ODE network, under the assumption of toric geometry. To accomplish fitting, it is necessary to account for initial conditions to predict *μ* and the exact eigendecomposition of Σ, but this may still be possible in the absence of simulation. Such an approach would not only take advantage of information unique to single-cells, but could also make it possible to parameterize models too complex for conventional fitting (fitting involves many rounds of simulation). Because it has explicit connections to CRNT, such a method could be used in conjunction with other recently developed applications of CRNT for data-constrained, ODE model selection [41–45]. This provides a principled way to choose among models with different components and topologies, a common goal of systems biology modeling projects.

One limitation of the network analysis approach described here is that identifying gaps in the eigenvalue spectrum is heuristic. Unfortunately, this is true of most other applications in which cutoffs in eigenvalue spectra must be identified. The relatively low dimensionality of FACS and CyCIF datasets further limits the applicability of principled approaches that are available, including those based on random matrix theory. However, for larger datasets it will potentially be possible to apply principled methods for identifying gaps that are statistically significant.

Cell regulatory networks are characterized by multistability and limit cycles. The relationship between our analysis and such network structures remains unclear and will require further theoretical analysis. Multistable, toric steady state sets exist, but there are many circumstances in which multistable states are not toric. Perhaps such non-toric sets can be approximated as toric in various limiting regimes, but even then, certain parts of the steady state set must necessarily be unstable to support multistability. For limit cycles, the expected geometry is not necessarily algebraic, although one could hope that the limit cycle is contained in an almost-toric manifold, so that our approach is still informative. Exploring these and other issues requires further development of the connection between the sc-data and CRNT.

The promise of the ESS approach is that it provides a potentially powerful but as-yet unexplored, geometric framework for linking features in sc-data to reaction networks. This is in parallel with recent geometrical analysis of CRNT, in which toric varieties have played a key role. Toric varieties have aided in characterizing the central CRNT concept of complex-balancing [25]. They have also enabled systematic determination of kinetic parameters that give rise to multistability for large classes of networks [46], including biologically relevant networks such as the MAPK pathway [47]. In the context of sc-data, we leverage toric geometry to study the reactions underlying cellular phenotypes without having to perform simulations, which can be difficult with sparse and complex data. Despite the fact that some network steady states do not necessarily conform to toric geometries, the CRNT framework as accessed through ESS is a closer approximation to the reality of biological networks than the statistical and dimensionality reduction approaches (clustering, tSNE etc.) that currently dominate data analysis. With further development of the ESS approach, it should be possible to use CRNT to formulate mechanistic hypotheses from data in the absence of simulation and then subject the hypotheses to empirical tests.

## Methods

### Sc-covariance matrix from complex-balanced reaction networks

For any complex-balanced reaction network *G*, the steady state set *E* is log-linear, so as *t* → ∞, each 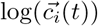 approaches a linear subset *V* = log(*E*). Therefore, in the limit, the sample covariance matrix is singular, and its singular eigenspace 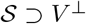, the orthogonal complement of *V*. Complex-balancing implies that *V*^⊥^ = *S*, the stoichiometric subspace of *G* [25], so 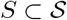.

The equality 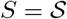 arises when the distribution of 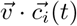, where *i* indexes over all cells, at *t* = 0 has non-zero variance 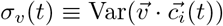 for all 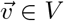. Splitting the chemical concentration space into vector-additive cosets of the stoichiometric subspace *S*, all trajectories of a complex-balanced system are forward-invariant within each coset. By orthogonality of *V* to *S*, *σ_v_*(*t*) is time-independent. If *σ_v_* is non-zero for all *v*, then the variance in log-concentration space of 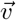, given by 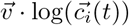 cannot be zero. Therefore, 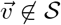, so 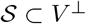. Together with the previous inclusion, 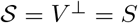, as *t* → ∞.

There is a unique steady state point *c_e_* in each coset of a complex-balanced reaction network [24], which has an exponentially stable neighborhood. The Global Attractor Conjecture, for which a proof was announced [48] but has not yet been peer reviewed, suggests that all complex-balanced reaction networks are globally asymptotically stable (relative to a stoichiometric coset), so for sufficiently long *T*, any finite collection of trajectories uniformly enter their exponentially stable neighborhoods. After this time *T*, any two trajectories *c*_1_(*t*) and *c*_2_(*t*) on the same coset obey

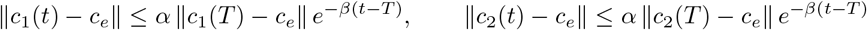

and so by the triangle inequality,

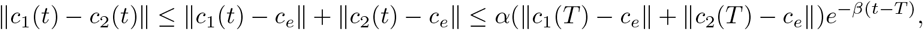

which gives a monotonically decreasing, upper bound on the distance between any pair of trajectories. As distance between all trajectories decreases, their variance decreases. As long as the steady state set has no boundary states (nontrivial equilibria with zero concentration for some chemical species), the logarithm of concentrations will also have a uniform upper-bound which decreases with time, after some time *T*′.

While the variance along the orthogonal complement of *V* decreases to 0, the variance along *V* remains non-zero, so the eigenvalues of the covariance matrix separate into those that are non-zero (large) and those approaching zero (small). Assuming no further network structure, at sufficiently large times, we consider the distribution of trajectories as a spiked population model in which the small eigenvalues follow a Marchenko-Pastur density with rescaled support [30], while the large eigenvalues lie outside its support. This forms a gap.

Furthermore, if the reaction network has a subnetwork with separably faster rates of convergence than the entire network, additional gaps may occur. In the case of detailed-balance reaction networks, this follows from a singular perturbation approach: Giovangigli et al. showed that for separably fast and slow reactions in such networks, the critical steady-state manifold is equivalent to the steady-state manifold of a network containing only the fast reactions [32]. Therefore, trajectories first converge to *V*_fast_ ⊃ *V*, leading to a gap between the fast, small eigenvalues and the slow, larger eigenvalues.

### Log-linearity despite unobservables

Given a *D*-dimensional log-linear set in ℝ^*N*^, parameterize it by a *D*-dimensional vector of parameters 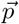 and *D* corresponding column vectors 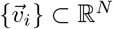 that span the set s.t.:

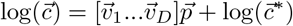

for some point 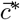 in the set that determines the translation of the affine set. For 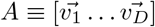, the orthogonal complement *S_N_* is the null space of *A^T^*.

Split the matrix *A* horizontally into two matrices, *A*_obs_, and *A*_unobs_, corresponding to *n* observed chemical species and *N* – *n* unobserved species. The vector whose coordinates are the observed species, 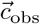, are parameterized by

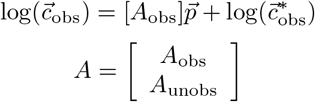

where the chemical species are rearranged for convenience, without loss of generality. Therefore, provided *n* > *D*, the data still lies in a nontrivial, log-linear set.

Now we show that the orthogonal complement of 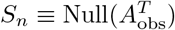, is a meaningful subspace:

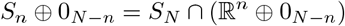

where 0_*N–n*_ is the zero-vector in the (*N – n*)-dimensional unobserved space.

#### Proof

For the forward inclusion, observe that *S_n_* ⊂ ℝ^*n*^, and simultaneously that *S_n_* ⊕ 0_*N–n*_ ⊂ *S_N_*, because for all 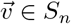,

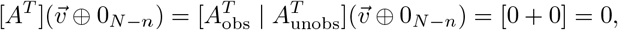

since by definition of 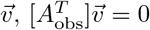.

For the reverse inclusion, a vector in (ℝ^*n*^ ⊕ 0_*N–n*_) takes the form 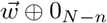, and being in *S_N_* indicates that

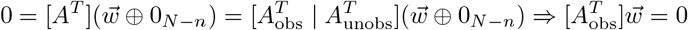

and therefore 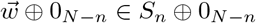.

### Simulation of random networks

ODE simulations of each reaction network were performed with the *ode*15*s* function in MATLAB. For each network, 300 initial conditions were chosen from a log-normal distribution with equal log-variance for all chemicals. Specific sampling distribution parameters are in Table 1.

**Table 1.** Simulation parameter distributions for randomized Complex-Balanced Networks (CB) and Gene Regulatory Networks (GRN)

| Parameter | Log-Mean | Log-Variance |
| --- | --- | --- |
| Concentration @t=0 (CB) | 4 | 4 |
| Kinetic Constants (CB) | 2.5, 3 | 0.05 |
| Concentration @t=0 (GRN) | 5 | 8 |
| Unbound Production Constants (GRN) | 1 | 1 |
| Bound Production Constants (GRN) | 3 | 3 |
| Protein Binding-Unbinding Constants (GRN) | 3 | 1 |

#### Complex-balanced networks

All complex-balanced networks were chosen to have *n* = 20 chemical species. Complex-balancing is defined via the digraph with nodes representing *complexes*, such as *A* + *B*, and edges representing reactions between complexes. A sufficient condition for complex-balancing is for a network to have *deficiency* equal to 0, and be *weakly reversible*. Deficiency *δ* is defined as

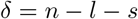

where *n* is the number of complexes, *l* is the number of weakly connected components, termed *linkage classes*, and *s* is the dimension of the stoichiometric subspace. Weak reversibility amounts to all nodes belonging to a strongly connected component.

Each random network was generated with all the nodes representing complexes containing either a single species, or any pair of species. Then, ~ 0.03% of the possible edges were stochastically chosen. The graph was then symmetrized by adding all the reverse edges, to ensure reversibility. Rate constants were randomly assigned from a log-normal distribution (see Table 1). Many such networks were generated, and only the ones with deficiency zero were simulated.

#### Gene regulatory networks

For the GRNs, for *n* genes, ~ 70% of the possible protein-bound genes 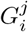 were chosen stochastically. Kinetic constants for each type of reaction were chosen from log-normal distributions whose log-mean and log-variance are shown in Table 1.

The GRN simulations were not complex-balanced, both because the particular arrangement of irreversible reactions violate weak reversibility, and because the deficiency of the networks were large, indicating a measure zero probability of being complex-balanced.

### Linear fluorescence assumption

The framework calls for analyzing chemical concentrations 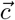. Both FACS and CyCIF data contain fluorescence intensity signals instead of the actual concentrations, but our method still applies if the *i*^th^ chemical species’ signal *I_i_* = *k_i_* · *c_i_* for some constant *k_i_* for the cells in one subpopulation. Assuming an excess of antibodies for both the experimental setup of the FACS and CyCIF data, this is simply the requirement that detection is in the linear regime.

The method still works because the *k_i_*’s would only result in a shift of the affine subspace *V*. For any log-linear constraint on the 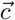

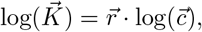

the observed constraint in terms of 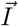 is

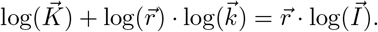

### Gaussian mixture modeling

Both FACS and CyCIF data were fit with Gaussian mixture models (GMM) to match visible clusters. Cells from any single treatment condition were fit with *k* components, with *k* chosen based on abrupt decreases in the incremental likelihood gain for additional components, while also preventing the splitting of visible clusters in the data.

GMMs were fit using the *fitgmdist* function in MATLAB 2016b, allowing the Expectation Maximization algorithm to run to convergence for at least 20k different initializations chosen by the *k*-means++ algorithm.

### Null distribution of row reduced vector entries

Assuming sparse, random, linear constraints on a distribution, the covariance matrix would have singular eigenvectors whose span can be given by sparse vectors with random orientations. For either the FACS or CyCIF data, the null was given by row reducing s vectors whose entries were chosen uniformly between −0.5 and 0.5, and subsequently made sparse at random entries. The dimension of the constraints, s, was chosen to be similar to that selected for each datasets’ stoichiometric subspace, and sparsity was set equal to the percentage of the zero-centered peak of the data’s entry-distribution in a window between −0. 2 and 0. 2. Gaussian noise was added to the final row-reduced vector entries, with variance matching that of the zero-centered window. Both null distributions were generated by Gaussian kernel smoothing of 1000 such sets of s row reduced vectors’ entries.

### Confidence intervals for row-reduced vector entries

The 95% confidence intervals for the row-reduced vector entries in Fig 4 were calculated by bootstrapping for 1000 replicates. Each time, the original data was resampled with replacement, before fitting a GMM with the same number of components as for the original data. Then, the stoichiometric subspace was chosen with the same dimension. Finally, row reduction was performed with the same column and row rearrangements as was done in the original data, instead of using the complete pivoting algorithm, to keep entries consistent between replicates.

## S1 Appendix. Steady state analysis of 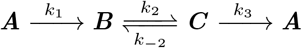 with unobservables

This network is deficiency zero, and weakly reversible, so it is complex-balanced. Additionally, there is one connected component of the reaction network, so [25] tells us that at steady state, for every pair of nodes *x, y*, the ratio of their values is

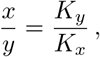

where *K_x_, K_y_* are defined based on the adjacency matrix of the network. Specifically, given the *n* × *n* weighted adjacency matrix *A* for the *n* nodes, where the kinetic constants *k_i_* are the weights, we construct the Laplacian *A_k_* of *A*, by subtracting the sum of each row from the corresponding entry of the diagonal. Then to determine *K_x_*, we remove the *x*’th column and row from *A_k_*, denote it *A_k\x_*, and calculate its determinant up to a sign. Rephrased as a formula:

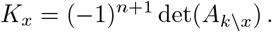

In our example, *K_B_* would be calculated as:

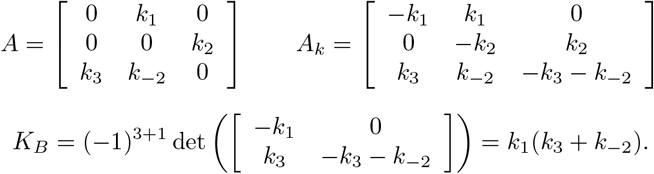

The quantities *K_A_, K_C_* can be calculated similarly, and their ratios are the constants in the steady state constraints describing (3).

Geometrically, the steady state set *E* specified by

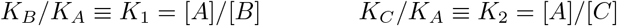

can be parameterized by *t* as a line:

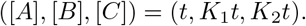

Supposing we only observe *A* and *B*, we are left with the line

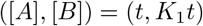

which, after taking the logarithms of each coordinate, becomes

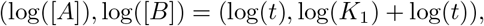

or in a different form, taking *T* = log(*t*):

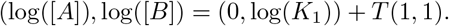

The orthogonal complement is spanned by (1, −1), since (1, −1) · (1, 1) = 0. This signifies the existence of a net balancing reaction between *A* and *B* in the full, unobserved network.

## S2 Appendix. Probability of manual gap choice producing observed vector entry distribution

In the FACS dataset, choosing a gap after *k* eigenvalues gives *k* eigenvectors each with *d* =11 entries. Some of these *dk* entries will be forced to 1 or 0 after row reduction, and so only (*d* − *k*)*k* entries are independent. In our data, we typically chose *k* = 7, leading to 28 independent entries. In the worst case, we could choose *k* = *d* − 1, leading to 10 free entries. Consider the observation that the distribution of entries was almost entirely negative or 0. For one condition, the probability of a given gap choice producing that asymmetry for a random set of vectors is at most 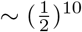, and the chance that at least one of the 10 gap choices gives the asymmetry is ~ 2 × 10^−3^.

However, suppose that we include sparseness of the vectors in our null to match the observed vectors’ sparseness, leading to a gap that includes *k* eigenvalues to specify ~ *k* nonzero, independent entries that are equally likely to be positive or negative. Most of our gaps had at least *k* > 5. The probability of *k* entries all being negative is 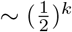, so that for a random dataset with sparse structure, the chance of any such gap existing is 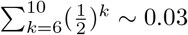. Pessimistically assuming that all 13 samples had identical structure, 0.03 is the probability that a random, sparse structure would even admit a gap choice that produces the observed asymmetry. This probability decreases further if we had lower sparsity, or if we account for the 13 samples having at least partially different structure due to the difference in perturbation conditions between them.

Additionally, our entry distribution showed peaks. The probability of the observed peaks’ prominence, *p*_peak_, is independent of the probability of asymmetry, although harder to estimate. Therefore, the probability that a dataset would even admit gap choices that produce the key features of our entry distribution is < (0.03)*p*_peak_.

**S1 Fig. Interaction Network from FACS** Given recombined, singular vectors for the treatment condition of activation with anti-CD3, anti-CD8 and inhibition of Protein Kinase C with G06976, we drew an edge between biomarkers if any vector entries had magnitude larger than 0.2.

## Acknowledgments

This work was supported by NCI grant U54-CA225088 and DARPA grant W911NF018-1-0124 (to PKS), AFOSR grant FA9550-14-1-0060 and NSF grant 1817936 (to EDS). SW was partially supported by NIH/NIGMS T32 GM008313.

## References

1. Bendall SC, Nolan GP. From single cells to deep phenotypes in cancer. Nature Biotechnology. 2012;30(7):639–647. doi:10.1038/nbt.2283.

2. Galli E, Friebel E, Ingelfinger F, Unger S, Núñez NG, Becher B. The end of omics? High dimensional single cell analysis in precision medicine. European Journal of Immunology. 2019;49(2):212–220. doi:10.1002/eji.201847758.

3. Nawy T. Single-cell sequencing. Nature Methods. 2014;11(1):18–18. doi:10.1038/nmeth.2771.

4. Perez OD, Nolan GP. Simultaneous measurement of multiple active kinase states using polychromatic flow cytometry. Nature Biotechnology. 2002;20(2):155–162. doi:10.1038/nbt0202-155.

5. Lin JR, Fallahi-Sichani M, Sorger PK. Highly multiplexed imaging of single cells using a high-throughput cyclic immunofluorescence method. Nature Communications. 2015;6(1). doi:10.1038/ncomms9390.

6. Lin JR, Izar B, Wang S, Yapp C, Mei S, Shah PM, et al. Highly multiplexed immunofluorescence imaging of human tissues and tumors using t-CyCIF and conventional optical microscopes. eLife. 2018;7. doi:10.7554/elife.31657.

7. Giesen C, Wang HAO, Schapiro D, Zivanovic N, Jacobs A, Hattendorf B, et al. Highly multiplexed imaging of tumor tissues with subcellular resolution by mass cytometry. Nature Methods. 2014;11(4):417–422. doi:10.1038/nmeth.2869.

8. Lubeck E, Cai L. Single-cell systems biology by super-resolution imaging and combinatorial labeling. Nature Methods. 2012;9(7):743–748. doi:10.1038/nmeth.2069.

9. Yuan GC, Cai L, Elowitz M, Enver T, Fan G, Guo G, et al. Challenges and emerging directions in single-cell analysis. Genome Biology. 2017;18(1). doi:10.1186/s13059-017-1218-y.

10. Wagner DE, Weinreb C, Collins ZM, Briggs JA, Megason SG, Klein AM. Single-cell mapping of gene expression landscapes and lineage in the zebrafish embryo. Science. 2018;360(6392):981–987. doi:10.1126/science.aar4362.

11. Cai L, Friedman N, Xie XS. Stochastic protein expression in individual cells at the single molecule level. Nature. 2006;440(7082):358–362. doi:10.1038/nature04599.

12. Shin YS, Remacle F, Fan R, Hwang K, Wei W, Ahmad H, et al. Protein Signaling Networks from Single Cell Fluctuations and Information Theory Profiling. Biophysical Journal. 2011;100(10):2378–2386. doi:10.1016/j.bpj.2011.04.025.

13. Dunlop MJ, Cox RS, Levine JH, Murray RM, Elowitz MB. Regulatory activity revealed by dynamic correlations in gene expression noise. Nature Genetics. 2008;40(12):1493–1498. doi:10.1038/ng.281.

14. van der Maaten L, Hinton G. Visualizing Data using t-SNE. Journal of Machine Learning Research. 2008;9:2579–2605.

15. Botsch M, Pajarola R, Singh G, Memoli F, Carlsson G. Topological methods for the analysis of high dimensional data sets and 3D object recognition. Eurographics Symposium on Point-Based Graphics. 2007; p. 91–100.

16. Sachs K. Causal Protein-Signaling Networks Derived from Multiparameter Single-Cell Data. Science. 2005;308(5721):523–529. doi:10.1126/science.1105809.

17. Satija R, Farrell JA, Gennert D, Schier AF, Regev A. Spatial reconstruction of single-cell gene expression data. Nature Biotechnology. 2015;33(5):495–502. doi:10.1038/nbt.3192.

18. Waage P, Gulberg CM. Studies concerning affinity. Journal of Chemical Education. 1986;63(12):1044. doi:10.1021/ed063p1044.

19. Johnson KA, Goody RS. The Original Michaelis Constant: Translation of the 1913 Michaelis–Menten Paper. Biochemistry. 2011;50(39):8264–8269. doi:10.1021/bi201284u.

20. Gesztelyi R, Zsuga J, Kemeny-Beke A, Varga B, Juhasz B, Tosaki A. The Hill equation and the origin of quantitative pharmacology. Archive for History of Exact Sciences. 2012;66(4):427–438. doi:10.1007/s00407-012-0098-5.

21. Chen WW, Niepel M, Sorger PK. Classic and contemporary approaches to modeling biochemical reactions. Genes & Development. 2010;24(17):1861–1875. doi:10.1101/gad.1945410.

22. Kazakiewicz D, Karr JR, Langner KM, Plewczynski D. A combined systems and structural modeling approach repositions antibiotics for Mycoplasma genitalium. Computational Biology and Chemistry. 2015;59:91–97. doi:10.1016/j.compbiolchem.2015.07.007.

23. Karr JR, Sanghvi JC, Macklin DN, Gutschow MV, Jacobs JM, Bolival B, et al. A Whole-Cell Computational Model Predicts Phenotype from Genotype. Cell. 2012;150(2):389–401. doi:10.1016/j.cell.2012.05.044.

24. Horn F, Jackson R. General mass action kinetics. Archive for Rational Mechanics and Analysis. 1972;47(2). doi:10.1007/bf00251225.

25. Craciun G, Dickenstein A, Shiu A, Sturmfels B. Toric dynamical systems. Journal of Symbolic Computation. 2009;44(11):1551–1565. doi:10.1016/j.jsc.2008.08.006.

26. Joshi B, Shiu A. A Survey of Methods for Deciding Whether a Reaction Network is Multistationary. Mathematical Modelling of Natural Phenomena. 2015;10(5):47–67. doi:10.1051/mmnp/201510504.

27. Tolman RC. The Principles of Statistical Mechanics (Dover Books on Physics). Dover Publications; 2010.

28. Pérez-Millán M, Dickenstein A, Shiu A, Conradi C. Chemical Reaction Systems with Toric Steady States. Bulletin of Mathematical Biology. 2011;74(5):1027–1065. doi:10.1007/s11538-011-9685-x.

29. Ringnér M. What is principal component analysis? Nature Biotechnology. 2008;26(3):303–304. doi:10.1038/nbt0308-303.

30. Baik J, Silverstein JW. Eigenvalues of large sample covariance matrices of spiked population models. Journal of Multivariate Analysis. 2006;97(6):1382–1408. doi:10.1016/j.jmva.2005.08.003.

31. Verhulst F. Singular perturbation methods for slow–fast dynamics. Nonlinear Dynamics. 2007;50(4):747–753. doi:10.1007/s11071-007-9236-z.

32. Giovangigli V, Massot M. Entropic structure of multicomponent reactive flows with partial equilibrium reduced chemistry. Mathematical Methods in the Applied Sciences. 2004;27(7):739–768. doi:10.1002/mma.429.

33. Bjorck A, Golub GH. Numerical Methods for Computing Angles Between Linear Subspaces. Mathematics of Computation. 1973;27(123):579. doi:10.2307/2005662.

34. Ferré L. Selection of components in principal component analysis: A comparison of methods. Computational Statistics & Data Analysis. 1995;19(6):669–682. doi:10.1016/0167-9473(94)00020-j.

35. Cangelosi R, Goriely A. Component retention in principal component analysis with application to cDNA microarray data. Biology Direct. 2007;2(1):2. doi:10.1186/1745-6150-2-2.

36. Golub GH, Loan CFV. Matrix Computations (Johns Hopkins Studies in Mathematical Sciences)(3rd Edition). Johns Hopkins University Press; 1996.

37. Cheng K, Hao M. Mammalian Target of Rapamycin (mTOR) Regulates Transforming Growth Factor-*β*1 (TGF-*β*1)-Induced Epithelial-Mesenchymal Transition via Decreased Pyruvate Kinase M2 (PKM2) Expression in Cervical Cancer Cells. Medical Science Monitor. 2017;23:2017–2028. doi:10.12659/msm.901542.

38. Qiu L, Zhang Y, Li CK. Unitarily Invariant Metrics on the Grassmann Space. SIAM Journal on Matrix Analysis and Applications. 2005;27(2):507–531. doi:10.1137/040607605.

39. Sun SY. mTOR kinase inhibitors as potential cancer therapeutic drugs. Cancer Letters. 2013;340(1):1–8. doi:10.1016/j.canlet.2013.06.017.

40. Fisher RA. The Use of Multiple Measurements in Taxonomic Problems. Annals of Eugenics. 1936;7(2):179–188. doi:10.1111/j.1469-1809.1936.tb02137.x.

41. Adamer MF, Helmer M. Complexity of model testing for dynamical systems with toric steady states. Advances in Applied Mathematics. 2019;110:42–75. doi:10.1016/j.aam.2019.06.001.

42. Craciun G, Kim J, Pantea C, Rempala GA. Statistical Model for Biochemical Network Inference. Communications in Statistics - Simulation and Computation. 2013;42(1):121–137. doi:10.1080/03610918.2011.633200.

43. Gross E, Harrington HA, Rosen Z, Sturmfels B. Algebraic Systems Biology: A Case Study for the Wnt Pathway. Bulletin of Mathematical Biology. 2015;78(1):21–51. doi:10.1007/s11538-015-0125-1.

44. Gross E, Davis B, Ho KL, Bates DJ, Harrington HA. Numerical algebraic geometry for model selection and its application to the life sciences. Journal of The Royal Society Interface. 2016;13(123):20160256. doi:10.1098/rsif.2016.0256.

45. Pantea C, Gupta A, Rawlings JB, Craciun G. The QSSA in Chemical Kinetics: As Taught and as Practiced. In: Discrete and Topological Models in Molecular Biology. Springer Berlin Heidelberg; 2013. p. 419–442. Available from: https://doi.org/10.1007/978-3-642-40193-0_20.

46. Dickenstein A, Pérez-Millán M, Shiu A, Tang X. Multistationarity in Structured Reaction Networks. Bulletin of Mathematical Biology. 2019;81(5):1527–1581. doi:10.1007/s11538-019-00572-6.

47. Perez-Millan M, Turjanski AG. MAPK’s networks and their capacity for multistationarity due to toric steady states. Mathematical Biosciences. 2015;262:125–137. doi:10.1016/j.mbs.2014.12.010.

48. Craciun G. Toric Differential Inclusions and a Proof of the Global Attractor Conjecture; arXiv preprint 1501.02860, 2015.

